## Supplementary Material for "A bifunctional snoRNA with separable activities in guiding rRNA 2’-O-methylation and scaffolding gametogenesis effectors"

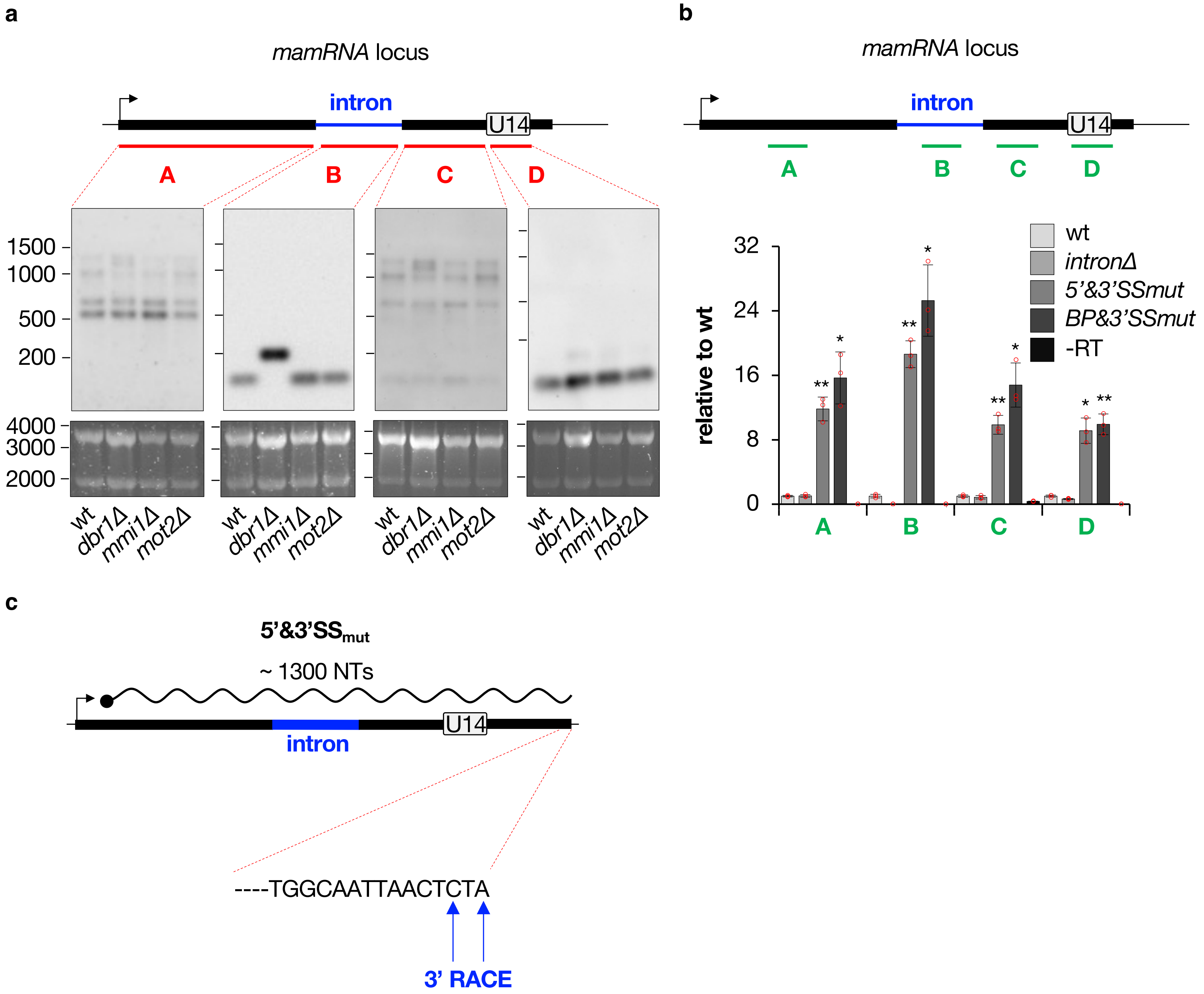


**Supplementary Fig. 1 (related to Fig. 1): *mamRNA* is produced from a *U14*-containing precursor and undergoes splicing. a**, Northern blots showing *mamRNA* and *U14* levels from total RNA samples, in the indicated genetic backgrounds. The position of the different probes used (A, B, C and D) are depicted on top. Ribosomal RNAs served as loading control (lower panels). **b**, RT-qPCR analyses of *mamRNA* and *U14* RNA levels in cells of the indicated genetic backgrounds (mean±SD; n=3; normalized to *act1+* and relative to wt). The position of the different amplicons analyzed (A, B, C, D) is indicated on top. Student's t-test (two-tailed) was used to calculate p-values (relative to wt). Individual data points are represented by red circles. **c**, 3' RACE analysis of the unspliced *mamRNA-U14* transcript in 5'&3'$SS_{mut}$ cells. The sequences of the 3' ends obtained from 2 independent clones are indicated by blue arrows.

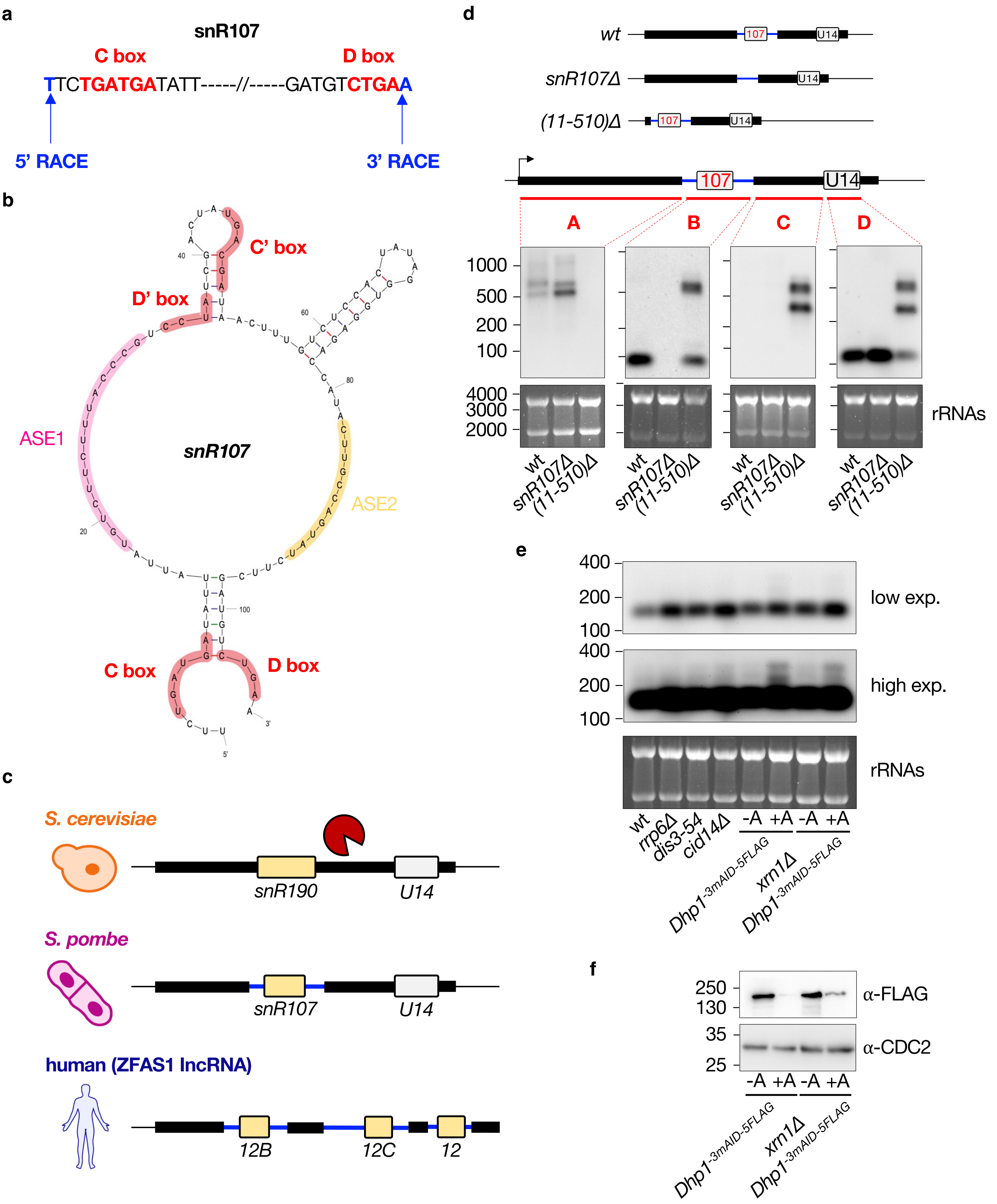


**Supplementary Fig. 2 (related to Fig. 2): *mamRNA* intron encodes a C/D-box snoRNA. a**, 5' and 3' RACE analyses of *snR107* in wild type cells. The sequences of the 5' and 3' ends were obtained from 1 and 2 independent clones respectively and are indicated by blue arrows. C and D boxes are labeled in red. **b**, Mfold prediction of *snR107* secondary structure. The terminal C and D as well as internal C' and D' boxes are highlighted in red. The ASE1 and ASE2, highlighted in magenta and orange respectively, were maintained single stranded. (continued)

**Supplementary Fig. 2 (continued): c**, Scheme depicting the genomic organization of *S. cerevisiae snR190*, *S. pombe snR107* and human *SNORD12/B/C*. Blue lines denote intronic sequences. **d**, Northern blots showing *mamRNA* and *snoU14* levels from total RNA samples, in the indicated genetic backgrounds represented on top. The position of the different probes used (A, B, C and D) are depicted. Ribosomal RNAs served as loading control. **e**, Northern blots showing *snR107* levels from total RNA samples, in the indicated genetic backgrounds. Ribosomal RNAs served as loading control. An auxin-inducible degron tagged version of Dhp1 was used for protein depletion. -A = no auxin; +A = auxin. **f**, Western blots showing the levels of an auxin-inducible degron tagged version of Dhp1 in the indicated genetic backgrounds in the absence (-A) or presence (+A) of auxin. An anti-CDC2 antibody was used as loading control.

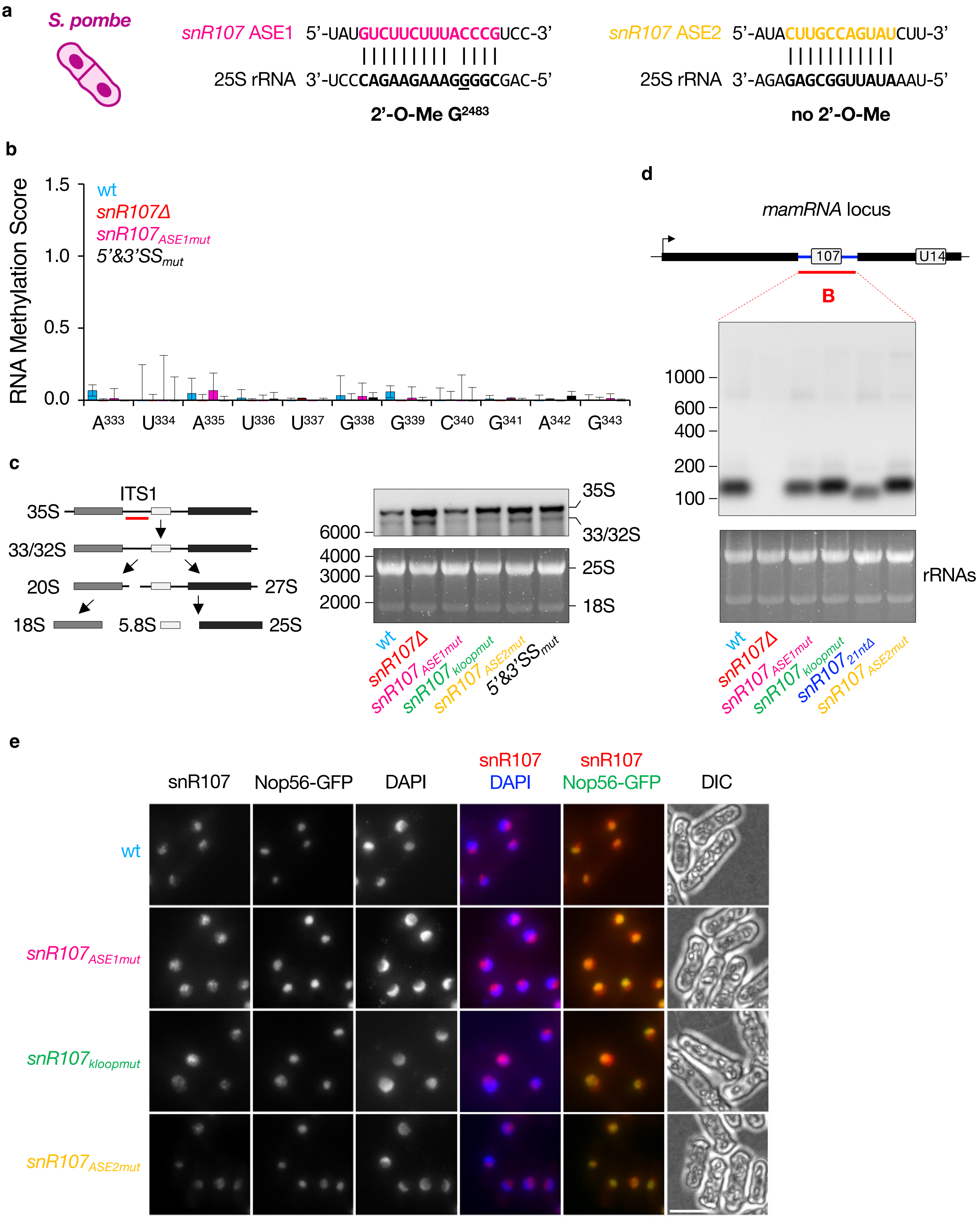


**Supplementary Fig. 3 (related to Fig. 3): *snR107* guides 25S rRNA 2’-O-methylation and is required for ribosome biogenesis. a**, Scheme showing the sequence complementarities between *snR107* ASE1 (magenta) or ASE2 (orange) and 25S rRNA (black). $G^{2483}$ is underlined. (continued)

**Supplementary Fig. 3 (continued): b**, RNA methylation scores (RMS) of the 25S rRNA residues 333 to 343 (complementary to *snR107* ASE2) as determined by RiboMethSeq in strains of the indicated genotypes ($mean_{(RMS-median\ all\ RMS)}$±SD; n=3)). **c**, Northern blot showing 35S and 33/32S pre-rRNA levels from total RNA samples, in the indicated genetic backgrounds. The position of the probe is indicated on the left as a red line. Ribosomal RNAs served as loading control. **d**, Northern blot showing *snR107* levels from total RNA samples, in the indicated genetic backgrounds. Ribosomal RNAs served as loading control. **e**, Representative images of *snR107* detected by smFISH in cells expressing GFP-tagged Nop56 in the indicated genetic backgrounds. DNA was stained with DAPI. Images are shown as Z-projections. Scale bar, 5 µm.

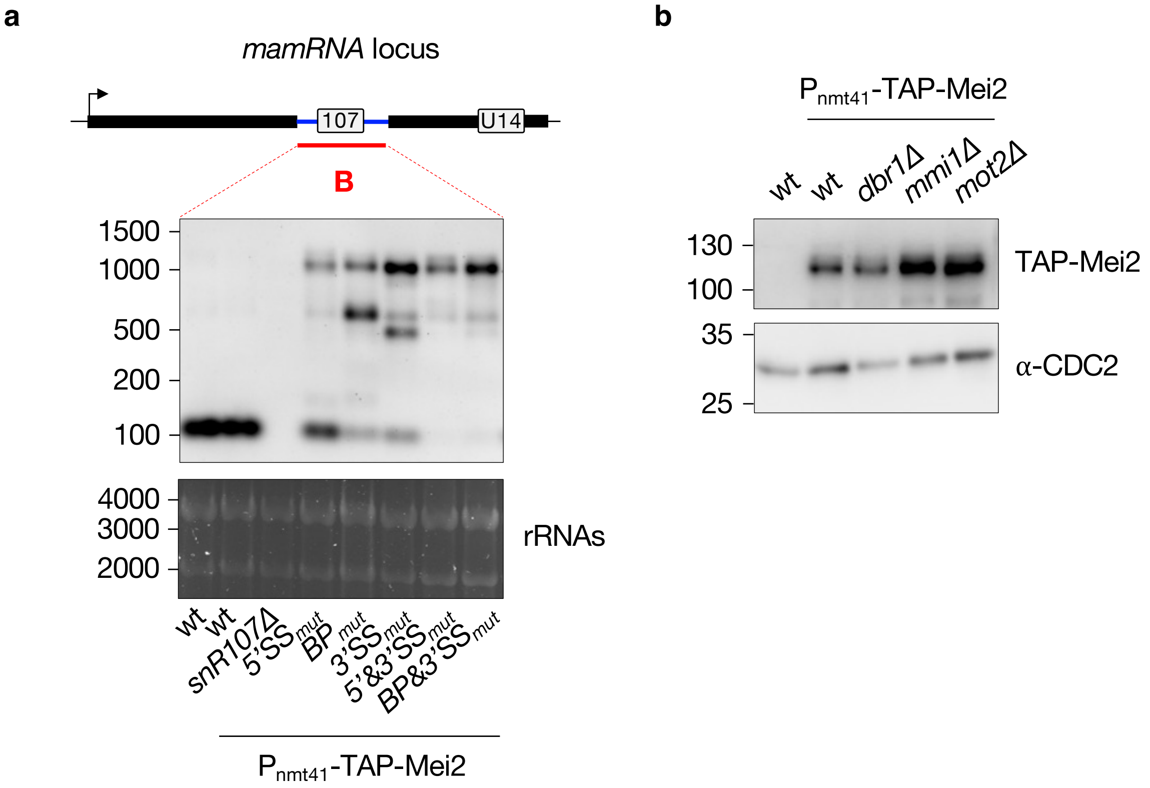


**Supplementary Fig. 4 (related to Fig. 4): *snR107* mediates the Mmi1-Mei2 mutual control. a**, Northern blot showing *snR107* levels from total RNA samples, in the indicated genetic backgrounds. Ribosomal RNAs served as loading control. **b**, Western blot showing the levels of TAP-tagged Mei2 in cells of the indicated genetic backgrounds. An anti-CDC2 antibody was used as loading control.

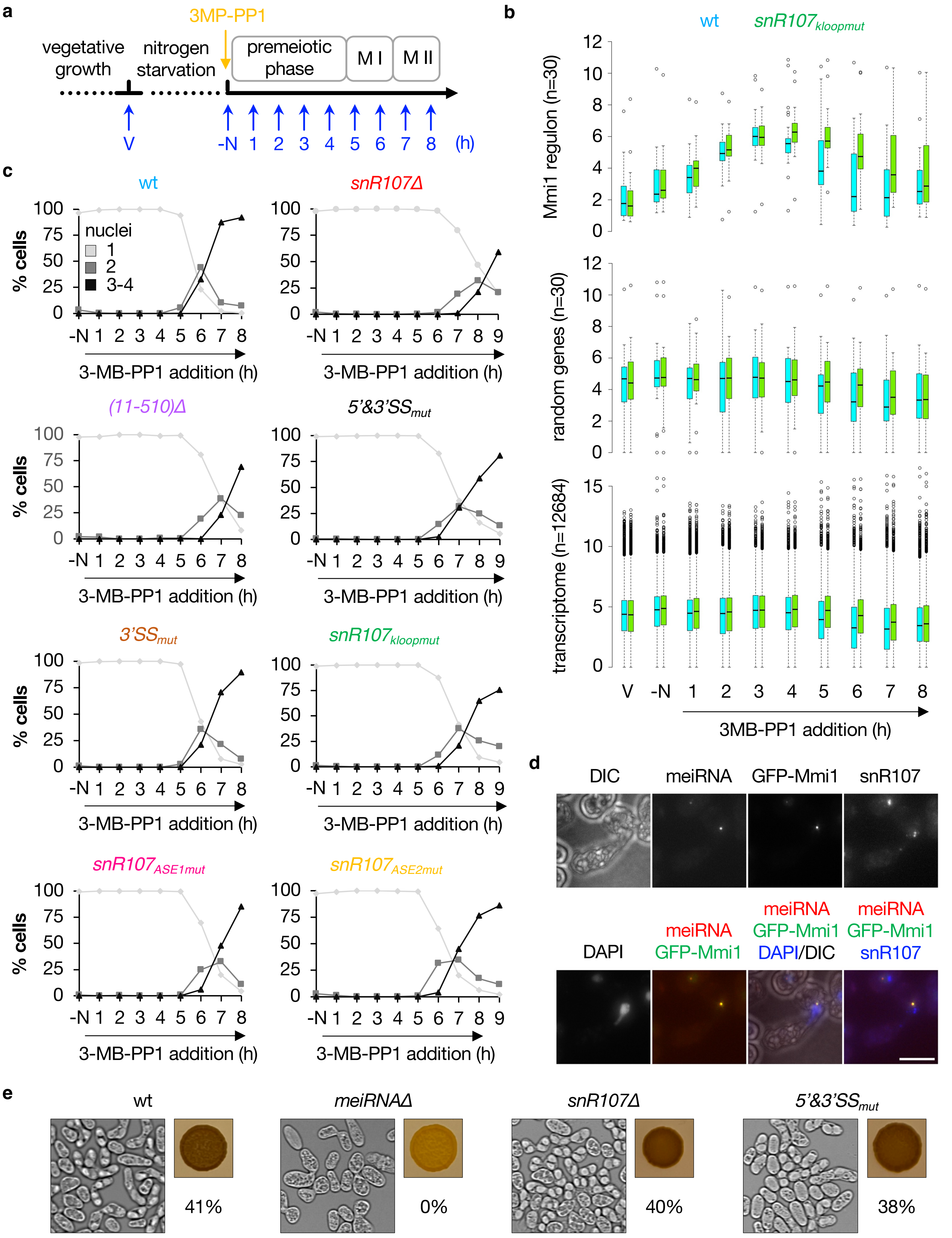


**Supplementary Fig. 5 (related to Fig. 5): *snR107* restricts the amplitude and timing of meiotic gene expression.**

**Supplementary Fig. 5 (related to Fig. 5): *snR107* restricts the amplitude and timing of meiotic gene expression.** **a**, Scheme illustrating the experimental strategy for the induction of meiosis. Following synchronization in G1 by nitrogen starvation (-N), meiosis was induced by adding 3-MB-PP1 to the culture medium and cells were harvested every hour as indicated. V = vegetative cells; -N = nitrogen-starved cells. **b**, Comparison of wild type and $snR107_{kloopmut}$ poly(A)+ transcriptomes by RNA-seq following induction of meiosis (n=2). Box plots show the relative expression (log2) of 30 Mmi1 targets[10] (upper panel), 30 random transcripts (middle panel) and the entire transcriptome (n=12684 RNAs). Box center lines represent the median, box limits represent the upper and lower quartiles, whiskers define the 1.5x interquartile range and individual points correspond to outliers. **c**, Meiotic time course in cells of the indicated genotypes. Hoechst-stained nuclei were counted at the indicated time points. Values represent the percentages of cells with one nucleus (diamonds), two nuclei (squares), three or four nuclei (triangles) from two independent experiments. **d**, Representative images of *meiRNA* and *snR107* detected by smFISH in wild type meiotic cells (horsetail stage of meiotic prophase I) expressing GFP-tagged Mmi1. DNA was stained with DAPI. Images are shown as Z-projections. Scale bar, 5 µm. **e**, Mating/sporulation efficiencies of the indicated homothallic strains (% tetrads), as determined by iodine staining and live cell imaging.

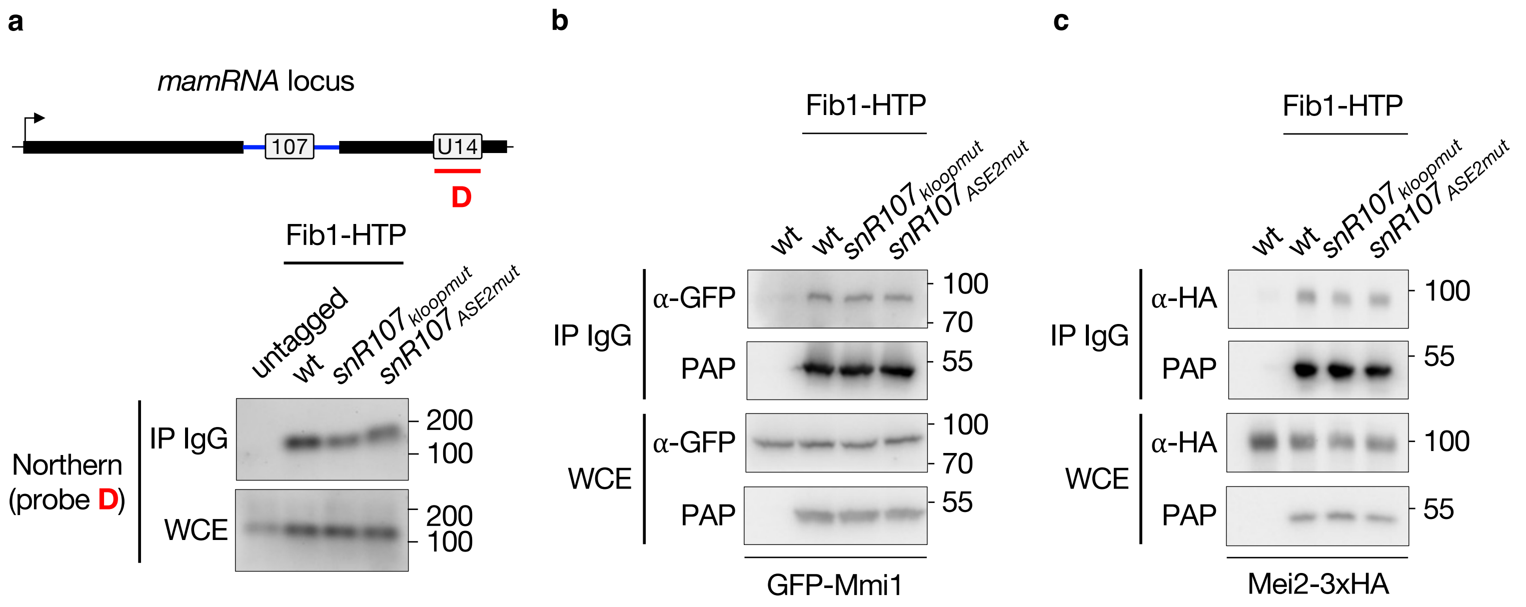


**Supplementary Fig. 6 (related to Fig. 6): Mmi1 and Mei2 associate with *snR107* and the 2'-O-methyltransferase Fib1. a**, Northern blot showing the fraction of *U14* (probe D) immunoprecipitated with HTP-tagged Fib1 in cells of the indicated genetic backgrounds. An untagged strain was used as negative control. IP = Immunoprecipitate; WCE = Whole Cell Extract. **b** and **c**, Western blots showing that GFP-tagged Mmi1 (**b**) and 3xHA-tagged Mei2 (**c**) co-immunoprecipitate with HTP-tagged Fib1 in cells of the indicated genotypes. The Fib1-Mei2 coIP (**c**) was carried out upon meiosis induction, 3 hours post Pat1 inactivation. IP = Immunoprecipitate; WCE = Whole Cell Extract.

**Supplementary Table 1: *S. pombe* strains used in this study**

| Strain | Genotype | Source |
|---|---|---|
| PR040 | h90, *ura4-DS/E, ade6-M210, leu1-32, mat3M::ura4+* | lab stock |
| PR162 | h90, *ura4-D18, ade6-M210, leu1-32, mat3M(EcoRV)::ade6+* | lab stock |
| PR202 | mat3M::gfp+::*nat*$^{R}$*MX*, *leu1-32, ura4-DS/E,* cid14::*hph*$^{R}$*MX* | This study |
| PR207 | mat3M::gfp+::*nat*$^{R}$*MX*, *leu1-32, ura4-DS/E, his2+/his2-?,* dis3-54 | This study |
| PR486 | PR040, rrp6::*hph*$^{R}$*MX* | Simonetti et al., 2017 |
| PR525 | PR162, mot2::*nat*$^{R}$*MX* | Andric et al., 2021 |
| PR617 | PR40, meiRNA::*hph*$^{R}$*MX* | Andric et al., 2021 |
| PR675 | PR40, *kan*$^{R}$*MX::*P$_{nmt41}$-TAP-Mei2 | Simonetti et al., 2017 |
| PR720 | PR40, mot2::*nat*$^{R}$*MX*, *kan*$^{R}$*MX::*P$_{nmt41}$-TAP-Mei2 | Simonetti et al., 2017 |
| PR726 | PR40, mei4::*nat*$^{R}$*MX*, mmi1::*hph*$^{R}$*MX*, *kan*$^{R}$*MX::*P$_{nmt41}$-TAP-Mei2 | Simonetti et al., 2017 |
| PR1034 | PR40, mei2::*hph*$^{R}$*MX*, pREP41::*LEU2* | Andric et al., 2021 |
| PR1407 | PR40, mei2::*hph*$^{R}$*MX*, pREP41-TAP-Mei2::*LEU2* | Andric et al., 2021 |
| PR1408 | PR40, mei2::*hph*$^{R}$*MX*, pREP41-TAP-Mei2$^{F644A}$::*LEU2* | Andric et al., 2021 |
| PR1513 | M(h-), *leu1-32*, pat1$^{L95G}$::*kan*$^{R}$ | J. Ayté |
| PR1609 | PR1513, Mei2-HTP::*nat*$^{R}$*MX* | This study |
| PR1849 | PR1513, mot2::*nat*$^{R}$*MX* | This study |
| PR1889 | PR1513, mei4::*nat*$^{R}$*MX*, mmi1::*hygTK* | This study |
| PR1954 | PR1513, mamRNA-intronΔ | This study |
| PR1956 | PR1513, mamRNA$_{5'_splice_site_mutated}$ (5'SSmut) | This study |
| PR1957 | PR1513, mamRNA$_{3'_splice_site_mutated}$ (3'SSmut) | This study |
| PR1958 | PR1513, mamRNA$_{5'_and_3'_splice_sites_mutated}$ (5'&3'SSmut) | This study |
| PR1968 | PR1513, mamRNA$_{(11\text{-}510)\Delta}$ | This study |
| PR1975 | PR40, mamRNA$_{intron\Delta}$, *kan*$^{R}$*MX::*P$_{nmt41}$-TAP-Mei2 | This study |
| PR1977 | PR40, mamRNA$_{5'_splice_site_mutated}$ (5'SSmut), *kan*$^{R}$*MX::*P$_{nmt41}$-TAP-Mei2 | This study |
| PR1978 | PR40, mamRNA$_{3'_splice_site_mutated}$ (3'SSmut), *kan*$^{R}$*MX::*P$_{nmt41}$-TAP-Mei2 | This study |
| PR1979 | PR40, mamRNA$_{5'_and_3'_splice_sites_mutated}$ (5'&3'SSmut), *kan*$^{R}$*MX::*P$_{nmt41}$-TAP-Mei2 | This study |
| PR1980 | PR40, mamRNA$_{(11\text{-}510)\Delta}$, *kan*$^{R}$*MX::*P$_{nmt41}$-TAP-Mei2 | This study |
| PR1984 | PR40, mamRNA$_{(11\text{-}737)\Delta}$, *kan*$^{R}$*MX::*P$_{nmt41}$-TAP-Mei2 | This study |

| | | |
|---|---|---|
| PR1992 | PR40, $mamRNA_{branchpoint\_mutated}$ (BPmut), *$kan^{R}MX$::*$P_{nmt41}$-TAP-Mei2 | This study |
| PR2001 | PR40, $mamRNA_{5'\_and\_3'\_splice\_sites\_mutated}$ (5'&3'SSmut) | This study |
| PR2004 | PR1513, $P_{mmi1}$-PTH-Mmi1 | This study |
| PR2012 | PR40, dbr1*::hygTK*, *$kan^{R}MX$::*$P_{nmt41}$-TAP-Mei2 | This study |
| PR2013 | PR1513, dbr1*::hygTK* | This study |
| PR2032 | PR1513, Fib1-HTP*::$nat^{R}MX$* | This study |
| PR2033 | PR1513, Nop56-HTP*::$nat^{R}MX$* | This study |
| PR2034 | PR1513, Nop58-HTP*::$nat^{R}MX$* | This study |
| PR2059 | PR1513, Fib1-HTP*::$nat^{R}MX$*, Mei2-3xHA*::$hph^{R}MX$* | This study |
| PR2066 | PR1513, $mamRNA_{branchpoint\_mutated}$ (BPmut) | This study |
| PR2067 | PR1513, $mamRNA_{branchpoint+3'\_splice\_site\_mutated}$ (BP+3'SSmut) | This study |
| PR2071 | PR1513, Mei2-3xHA*::$hph^{R}MX$* | This study |
| PR2079 | PR40, $snR107_{ASE1mut}$, *$kan^{R}MX$::*$P_{nmt41}$-TAP-Mei2 | This study |
| PR2083 | PR40, snR107Δ, *$kan^{R}MX$::*$P_{nmt41}$-TAP-Mei2 | This study |
| PR2097 | PR40, $snR107_{21nt\Delta}$, *$kan^{R}MX$::*$P_{nmt41}$-TAP-Mei2 | This study |
| PR2099 | PR40, $snR107_{ASE2mut}$, *$kan^{R}MX$::*$P_{nmt41}$-TAP-Mei2 | This study |
| PR2101 | PR1513, snR107Δ | This study |
| PR2102 | PR40, $P_{mmi1}$-GFP-Mmi1 | This study |
| PR2106 | PR40, $snR107_{kloopmut}$, *$kan^{R}MX$::*$P_{nmt41}$-TAP-Mei2 | This study |
| PR2111 | PR40, $mamRNA_{branchpoint+3'\_splice\_site\_mutated}$ (BP+3'SSmut), *$kan^{R}MX$::*$P_{nmt41}$-TAP-Mei2 | This study |
| PR2116 | M(h-), *ura4-D18*, ade6::ade6+-$P_{adh15}$-skp1-AtTIR1-2NLS $P_{adh15}$-skp1-OsTIR1, Dhp1-3xmAID-5xFLAG*::$kan^{R}MX$* | YGRC (FY39932) |
| PR2117 | PR1513, $snR107_{kloopmut}$ | This study |
| PR2120 | PR1513, $snR107_{ASE1mut}$ | This study |
| PR2121 | PR1513, $snR107_{ASE2mut}$ | This study |
| PR2125 | PR1513, Snu13-HTP*::$nat^{R}MX$* | This study |
| PR2129 | PR1513, $snR107_{kloopmut}$, Mei2-HTP*::$nat^{R}MX$* | This study |
| PR2136 | PR1513, $snR107_{kloopmut}$, Fib1-HTP*::$nat^{R}MX$* | This study |
| PR2137 | PR1513, $snR107_{ASE2mut}$, Mei2-HTP*::$nat^{R}MX$* | This study |
| PR2141 | PR162, mot2*::$nat^{R}MX$*, snR107Δ | This study |
| PR2143 | PR162, mot2*::$nat^{R}MX$*, $snR107_{kloopmut}$ | This study |
| PR2144 | PR162, mot2*::$nat^{R}MX$*, $mamRNA_{5'\_and\_3'\_splice\_sites\_mutated}$ (5'&3'SSmut) | This study |
| PR2145 | PR162, mot2*::$nat^{R}MX$*, $snR107_{ASE1mut}$ | This study |
| PR2146 | PR162, mot2*::$nat^{R}MX$*, $snR107_{ASE2mut}$ | This study |
| PR2148 | M(h-), *ura4-D18*, ade6::ade6+-$P_{adh15}$-skp1-AtTIR1-2NLS $P_{adh15}$-skp1-OsTIR1, Dhp1-3xmAID-5xFLAG*::$kan^{R}MX$*, xrn1*::$nat^{R}MX$* | This study |
| PR2155 | PR1513, $snR107_{kloopmut}$, $P_{mmi1}$-PTH-Mmi1 | This study |

| | | |
|---|---|---|
| PR2156 | PR1513, snR107$_{ASE2mut}$, P$_{mmi1}$-PTH-Mmi1 | This study |
| PR2158 | PR1513, snR107$_{ASE2mut}$, Fib1-HTP*::nat*$^{R}$*MX* | This study |
| PR2159 | PR1513, snR107$_{kloopmut}$, Nop58-HTP*::nat*$^{R}$*MX* | This study |
| PR2160 | PR1513, snR107$_{ASE2mut}$, Nop58-HTP*::nat*$^{R}$*MX* | This study |
| PR2161 | PR162, Nop56-GFP*::kan*$^{R}$*MX* | This study |
| PR2166 | PR1513, snR107$_{kloopmut}$, Fib1-HTP*::nat*$^{R}$*MX*, Mei2-3xHA*::hph*$^{R}$*MX* | This study |
| PR2167 | PR1513, snR107$_{ASE2mut}$, Fib1-HTP*::nat*$^{R}$*MX*, Mei2-3xHA*::hph*$^{R}$*MX* | This study |
| PR2175 | PR40, mamRNA$_{(11\text{-}510)\Delta}$, P$_{mmi1}$-GFP-Mmi1 | This study |
| PR2176 | PR40, mamRNA$_{intron\Delta}$, P$_{mmi1}$-GFP-Mmi1 | This study |
| PR2177 | PR40, snR107Δ, P$_{mmi1}$-GFP-Mmi1 | This study |
| PR2178 | PR40, mamRNA$_{5'_and_3'_splice_sites_mutated}$ (5'&3'SSmut), P$_{mmi1}$-GFP-Mmi1 | This study |
| PR2181 | PR162, snR107$_{ASE2mut}$, Nop56-GFP*::kan*$^{R}$*MX* | This study |
| PR2188 | PR40, snR107Δ | This study |
| PR2190 | PR162, snR107$_{kloopmut}$, Nop56-GFP*::kan*$^{R}$*MX* | This study |
| PR2192 | PR40, Fib1-HTP*::nat*$^{R}$*MX*, P$_{mmi1}$-GFP-Mmi1 | This study |
| PR2193 | PR40, snR107$_{kloopmut}$, Fib1-HTP*::nat*$^{R}$*MX*, P$_{mmi1}$-GFP-Mmi1 | This study |
| PR2194 | PR40, snR107$_{ASE2mut}$, Fib1-HTP*::nat*$^{R}$*MX*, P$_{mmi1}$-GFP-Mmi1 | This study |
| PR2196 | PR162, mamRNA$_{5'_and_3'_splice_sites_mutated}$, Nop56-GFP*::kan*$^{R}$*MX* | This study |
| PR2205 | PR1513, mei4*::nat*$^{R}$*MX*, P$_{mmi1}$-PTH-Mmi1$^{Y352F}$ | This study |
| PR2206 | PR162, snR107$_{ASE1mut}$, Nop56-GFP*::kan*$^{R}$*MX* | This study |
| PR2220 | PR1513, snR107$_{21nt\Delta}$ | This study |

**Supplementary Table 2: Plasmids used in this study**

| Number | Vector | Purpose | Source |
|---|---|---|---|
| B3874 | pFA6a-kanMX6, amp$^R$, ori | gene deletion | Lab stock |
| B3877 | pFA6a-GFP(S65T)-kan$^R$MX6, amp$^R$, ori | C-terminal tagging | Lab stock |
| BHM1296 | pFA6a-nat$^R$MX6, amp$^R$, ori | gene deletion | Lab stock |
| BHM1524 | pRS316-3xHA-hph$^R$MX, amp$^R$, ori | C-terminal tagging | Lab stock |
| BMR006 | pRS316-hph$^R$MX-P$_{act1}$-Ers1(first 500 nucleotides), amp$^R$, ori, URA3, CEN | gene deletion (hph$^R$MX cassette) | Lab stock |
| BMR015 | pREP41 (P$_{nmt41}$), LEU2, amp$^R$, ori | plasmid-driven expression | Lab stock |
| BMR021 | pBS1539-HTP-nat$^R$MX, amp$^R$, ori | C-terminal HTP tagging | Lab stock |
| BMR193 | pREP41-TAP-Mei2 (P$_{nmt41}$), LEU2, amp$^R$, ori | plasmid-driven expression | Lab stock |
| BMR194 | pREP41-TAP-Mei2$^{F644A}$ (P$_{nmt41}$), LEU2, amp$^R$, ori | plasmid-driven expression | Lab stock |
| BMR208 | pFA6a-HyTkAX | gene deletion | Addgene, #73898 |

**Supplementary Table 3: Oligonucleotides used in this study**

| Number | Location | Sequence | Assay |
|---|---|---|---|
| P249 | *mei4+* fwd | 5'-TGGATCAGATCCGTGGAATC-3' | RT-qPCR |
| P250 | *mei4+* rev | 5'-AACGCTCGATTAGAAGGCAT-3' | |
| P253 | *act1+* fwd | 5'-AACCCTCAGCTTTGGGTCTT-3' | |
| P254 | *act1+* rev | 5'-TTTGCATACGATCGGCAATA-3' | |
| P325 | *ssm4+* fwd | 5'-ACACAGTTTACGGGATTCTA-3' | |
| P326 | *ssm4+* rev | 5'-GATTGTGATGAAAACTGGGT-3' | |
| P607 | *mcp5+* fwd | 5'-AGACGTATTCACCTTACCTC-3' | |
| P608 | *mcp5+* rev | 5'-GTTTCCCATCATGACATGTT-3' | |
| P1208 | *mamRNA 5'* fwd (amplicon A) | 5'-TGGGATTAAGTCATCTTGGTG-3' | |
| P1209 | *mamRNA 5'* rev (amplicon A) | 5'-GTCAAAAAGATACCACAGCA-3' | |
| P1392 | *U14* fwd (amplicon D) | 5'-CAGGTGATGAAATTTCCATTG-3' | |
| P1284 | *U14* rev (amplicon D) | 5'-CATCCAAAGGAAGGACTATG-3' | |
| P1859 | *between mamRNA intron and U14* fwd (amplicon C) | 5'-ACGTGTATGTTCTGTATGGTG-3' | |
| P1924 | *between mamRNA intron and U14* rev (amplicon C) | 5'-CCAAAAAGCAATACAGCACTATA TATG-3' | |
| P2117 | *snR107* fwd (amplicon B) | 5'-ATGTCTTCTTTACCCGTCCTATC-3' | |
| P2118 | *snR107* rev (amplicon B) | 5'-CATCGAAGATACTGGCAAGTATG-3' | |
| P1313 | *mamRNA 5'* fwd | 5'-TAGGATGGTTCTCCTTCAGG-3' | RT-PCR |
| P1284 | *U14* rev | 5'-CATCCAAAGGAAGGACTATG-3' | |
| P1517 | *mamRNA 5'* fwd (probe A) | 5'-TCTGAAGGAGCCACATCAA-3' | Northern (T7 promoter-driven RNA probe) |
| P2101 | *mamRNA 5'* rev (probe A) | 5'-TAATACGACTCACTATAGGGCAAA CAAGTTTAACAAGGCAC-3' | |
| P2103 | *snR107* fwd (probe B) | 5'-GTATGTTCGCCGTATGCTG-3' | |
| P2104 | *snR107* rev (probe B) | 5'-TAATACGACTCACTATAGGGGAAG TTAGTACACTGAAAGATTATACAC-3' | |
| P1859 | *between mamRNA intron and U14* fwd (probe C) | 5'-ACGTGTATGTTCTGTATGGTG-3' | |

| | | | |
|---|---|---|---|
| P2211 | *between mamRNA intron and U14* rev (probe C) | 5'-TAATACGACTCACTATAGGGAACC TTCGCCGAACAATCCA-3' | |
| P1392 | *U14* fwd (probeD) | 5'-CAGGTGATGAAATTTCCATTG-3' | |
| P1449 | *U14* rev (probeD) | 5'-TAATACGACTCACTATAGGGCGTCA GAAAACACCAGCTGC-3' | |
| P866 | *meiRNA* fwd | 5'-CGGAATATGCATGCAAGAT-3' | |
| P1423 | *meiRNA* rev | 5'-TAATACGACTCACTATAGGGGTTTC AACAATAGTTCAGGTACAGTCG-3' | |
| P2355 | *25S rRNA 3'* fwd | 5'-TGATGAAGTGTCGTCGCAATG-3' | |
| P2356 | *25S rRNA 3'* rev | 5'-TAATACGACTCACTATAGGGGATAC AGGAGAGTAGGAACAAGTC-3' | |
| P2362 | *ITS1* fwd | 5'-CTCTTATCCATTTACCTTTCTGTG-3' | |
| P2363 | *ITS1* rev | 5'-TAATACGACTCACTATAGGGAGTAA TGATATGCTTGGCATGC-3' | |
| P2251 | *snR107* rev | 5'-TACAAGGGATCCCATCGAAGATAC TGGCAAGTATG-3' | 5' RACE (BamHI site) |
| P2230 | *snR107* fwd | 5'-GGACGGGTAAAGAAGACATAGCG ATATCATCAGAACAGCTTCG-3' | 3' RACE (NdeI site) |
| P2131 | *snR107* fwd #2 | 5'-ACGTTTCATATGACAACGAAGCTGT TCTGATG-3' | |
| P2375 | *downstream of U14* fwd | 5'-GTGTTTGGCTTGTAGTTGTAG-3' | |
| P2376 | *downstream of U14* fwd #2 | 5'-GATAAAGGTTGATTATCATGTTCTC C-3' | |
| P2357 | *25S rRNA RNA/DNA oligonucleotide* * | 5'-mUmUmUmUCCCCmGmCmUmGmAmU mUmUmUmGmCmC-3' | RNAseH cleavage |

* mN: 2'-O-methylated nucleotide

**Supplementary Table 4: smFISH probes used in this study**

| Target RNA | Probe | Sequence |
|---|---|---|
| *mamRNA 5'* (Quasar 670) | #1 | 5'-GTGGCTCCTTCAGACTCATG-3' |
| | #2 | 5'-TGGCGTCCTAGCATATTAAT-3' |
| | #3 | 5'-TAAACCTCTTGTCTCCACTA-3' |
| | #4 | 5'-GCCTTTCAACATCAGATAAT-3' |
| | #5 | 5'-GCCATTTACGATAAGAACCT-3' |
| | #6 | 5'-AAGAGCCTGAAGGAGAACCA-3' |
| | #7 | 5'-ACACAGATACAGGCGAACGC-3' |
| | #8 | 5'-ATCAGACTAGAATCTCTCCA-3' |
| | #9 | 5'-GCAACCATACACGACAGACA-3' |
| | #10 | 5'-ACCAAGATGACTTAATCCCA-3' |
| | #11 | 5'-CCAACACAGTACCAATTACT-3' |
| | #12 | 5'-AGGAATATGGTAGGTTTTTC-3' |
| | #13 | 5'-AAAGATACCACAGCAAGCGC-3' |
| | #14 | 5'-ACTAAGCTTTAACACCCAGT-3' |
| | #15 | 5'-AGTATGCGTACGGGATTTAC-3' |
| | #16 | 5'-TTCAGTTTGAGTTCAAGGTT-3' |
| *snR107* (Quasar 570) | #1 | 5'-CATAGTCGATAGGACGGGTA-3' |
| | #2 | 5'-TCCACCTATAGTGGAGACAA-3' |
| | #3 | 5'-AGACATCGAAGATACTGGCA-3' |
| | #4 | 5'-ATAGGACGGGTAAAGAAGAC-3' |
| | #5 | 5'-GTGGAGACAAAGTTATCGTC-3' |
| | #6 | 5'-GCAAGTATGGTCTCCACCTA-3' |
| | #7 | 5'-TCGTCATAGTCGATAGGACG-3' |
| | #8 | 5'-TACTGGCAAGTATGGTCTCC-3' |
| *meiRNA** (Quasar 670) | #1 | 5’-ATACCCACTAAGTCTGTTTA-3’ |
| | #2 | 5’-CGGCAGAAGATTGACCAACA-3’ |
| | #3 | 5’-GCATATTCCGTCTTACAATA-3’ |
| | #4 | 5’-ACCAACTAAAGCGATCTTGC-3’ |
| | #5 | 5’-GACCATTTCAAAATGTTGCA-3’ |
| | #6 | 5’-TACCGAATCCAGCTTTTTGA-3’ |
| | #7 | 5’-CAGAGCTTAGAAGACAAGGT-3’ |
| | #8 | 5’-TAACTGGACCCCATCAAGAA-3’ |
| | #9 | 5’-TAAACCAACTTGGGGGTTGG-3’ |
| | #10 | 5’-TCTAAGCTACTATTCATCCA-3’ |
| | #11 | 5’-AGTAGATTCCATCAGTCATA-3’ |
| | #12 | 5’-TGCAGCCAAAAAGTGTACCA-3’ |
| | #13 | 5’-CATTGTAAGTGCTTTCAAGG-3’ |
| | #14 | 5’-TTCAGTCATTCGCAAAGTTT-3’ |

| | | |
|---|---|---|
| | #15 | 5’-AGTCGTTTTATTTCTTTTCT-3’ |
| | #16 | 5’-GTTTCAACAATAGTTCAGGT-3’ |
| | #17 | 5’-TCTGTTTCAGGAATACGTTT-3’ |
| | #18 | 5’-TGTTTCGCATCAAACTTTCA-3’ |
| | #19 | 5’-GCGTTTAAACAAACTGCGGG-3’ |
| | #20 | 5’-TGGTTTCAGCACGTTTTCAA-3’ |
| | #21 | 5’-TTGGTTTGCAGGGTTTAACG-3’ |
| | #22 | 5’-CTTGCTGTGGTTATTGTTTA-3’ |

* same probe set as in Andric et al., 2021
